# Targeting human Plasmacytoid dendritic cells through BDCA2 prevents inflammation and fibrosis in xenotransplant mouse model of Scleroderma

**DOI:** 10.1101/2020.01.30.925073

**Authors:** Rebecca L. Ross, Clarissa Corinaldesi, Gemma Migneco, Ian Carr, Agne Antanaviciute, Antonio Carriero, Christopher Wasson, Ioanna Georgiou, Jörg H. W. Distler, Steve Holmes, Yasser M. El-Sherbiny, Clive S. McKimmie, Francesco Del Galdo

## Abstract

Plasmacytoid dendritic cells (pDC) have been implicated in the pathogenesis of Scleroderma (SSc) through their ability to infiltrate the skin and secrete interferons (IFN) and proinflammatory chemokines. Blood Dendritic Cells Antigen 2 (BDCA2) is an inhibitory type II C-type lectin expressed by human pDC. Here we determined the effects of BDCA2 internalisation on pDC mediated skin inflammation and fibrosis in human preclinical models of skin inflammation and fibrosis in vitro and in vivo. BDCA2 targeting reversed TLR-signalling induced transcriptome and differentiation of pDC and suppressed their ability to induce IFN response in organotypic 3D human skin cultures *in vitro*. *In vivo*, xenotransplantation of human pDC into immunocompromised mice (XenoSCID) significantly increased IFN induced responses to topical TLR7/9 agonist and separately enhanced the fibrotic response to bleomycin. Targeting of BDCA2 strongly suppressed both of these pathological responses ameliorating skin inflammation and fibrosis. Together, these preclinical data strongly support the notion that human pDC play a key role in immune driven skin fibrosis, which can be effectively blocked by targeting BDCA2.

## Introduction

Plasmacytoid dendritic cells (pDC) are bone marrow-derived myeloid cells specialized in the secretion of type I IFN [1, 2]. pDC activate inflammatory responses through TLR-mediated sensing of nucleic acid released from pathogen during infection or following cell death in autoimmunity. [3, 4]. Indeed, self-derived nucleic acids released from damaged tissues, apoptotic/necrotic cells or bound to autoantibodies, can be recognized by TLR7/8/9 and have been shown to induce pDC activation and IFN secretion [5–8].

Systemic sclerosis (SSc) is an immune-mediated inflammatory disease (IMID) characterized by vascular and tissue fibrosis, leading to diverse life-altering and life-threatening clinical manifestations [9]. pDC have been observed in affected skin of SSc patients and purified peripheral SSc pDC have been shown to spontaneously produce higher levels of type I IFN compared to healthy volunteers (HV) [10, 11]. Indeed, an elevated IFN gene signature in affected organs and in the blood is a common feature of severe disease in SSc. Accordingly, IFN inducible genes and those involved in activating an IFN response, such as CXCL4, CXCL9, CXCL10 or CXCL11 and their relative sera chemokines, have been identified as biomarkers for progression in SSc [11, 12]. Further, Brkic *et al.* have suggested that the same IFN signature is present before the onset of clinical fibrosis [13]. These observations collectively support the notion that type I IFNs may play an important role in the pathogenesis of SSc. Importantly, it is still not yet clear whether human pDC is the key cell type to drive the IFN response within the diseased skin and how this leads to skin inflammation and fibrosis.

pDC express multiple receptors that can signal to inhibit type I IFN secretion [2, 14]. One of these receptors is BDCA2 (CD303) [15]. BDCA2 is a type II transmembrane glycoprotein that belongs to the C-type lectin (CTLs) superfamily [16]. BDCA2 signals through an associated transmembrane adaptor, the FcϵRγ, which recruits the protein tyrosine kinase Syk, inducing protein tyrosine phosphorylation and calcium mobilization [17], which, in turn, interferes with TLR induced activation of pDC, inhibiting type I IFN secretion and other inflammatory mediators [15, 17–19]. For this reason, antibodies binding BDCA2 have been explored for their potential to block pDC activation in some IFN-associated autoimmune conditions, such as Systemic lupus erythematosus (SLE) [20, 21]. Recently, Ah Kioon and colleagues suggested that this could also be applicable to fibrotic disease such as SSc as mouse pDC enhance pathogenesis of bleomycin-induced skin fibrosis [10]. However, defining the efficacy of targeting BDCA2 during fibrosis is difficult in the mouse, as BDCA2 is only expressed in primates. Here we developed a fully-humanised monoclonal antibody with high affinity for BDCA2 and showed that BDCA2 targeting is sufficient to inhibit TLR induced human pDC activation *in vitro* and *ex vivo*. Further, we developed a xenotransplant model of human pDC in immunocompromised mice (XenoSCID) and defined the effects of pDC and BDCA2 targeting in TLR-induced skin inflammation and bleomycin-induced skin fibrosis. Our data clearly highlights the ability of TLR-activated human pDC to coordinate a skin IFN-response and results in skin inflammation and fibrosis, which is specifically inhibited by BDCA2 targeting.

## Results

### CBS004 specifically targets BDCA2 and blocks TLR9 induced human pDC IFN secretion

A monoclonal antibody targeted against BDCA2 (clone AC144) has previously been shown to suppress human pDC TLR-induced IFN type I secretion by interfering with the FcϵRγ-Syk signalling. We generated mouse monoclonal antibodies (mAb) against human BDCA2 and fully humanized the lead mAb. The affinity of the humanized antibody was measured by BIAcore, which exhibited 270-fold higher affinity for BDCA2 compared to AC144 control with a KD to <10 pM (limited by the BIAcore instrument detection limit) (Fig. 1A). Binding-competition assay by Enzyme-Linked Immunosorbent Assay (ELISA) determined that the affinity of CBS004 for human BDCA2 was 2-fold higher than AC144 control (Fig. 1A). *Ex vivo*, by using direct competition assays we showed that CBS004 and AC144 bind alternative epitopes as indicated by double staining of the pDC population gated within human peripheral blood mononuclear cells (PBMC) (LIN^−^ HLA^+^ CD123^+^ CD304^+^) (Fig. 1B, Fig. S1).

**Fig. 1.**
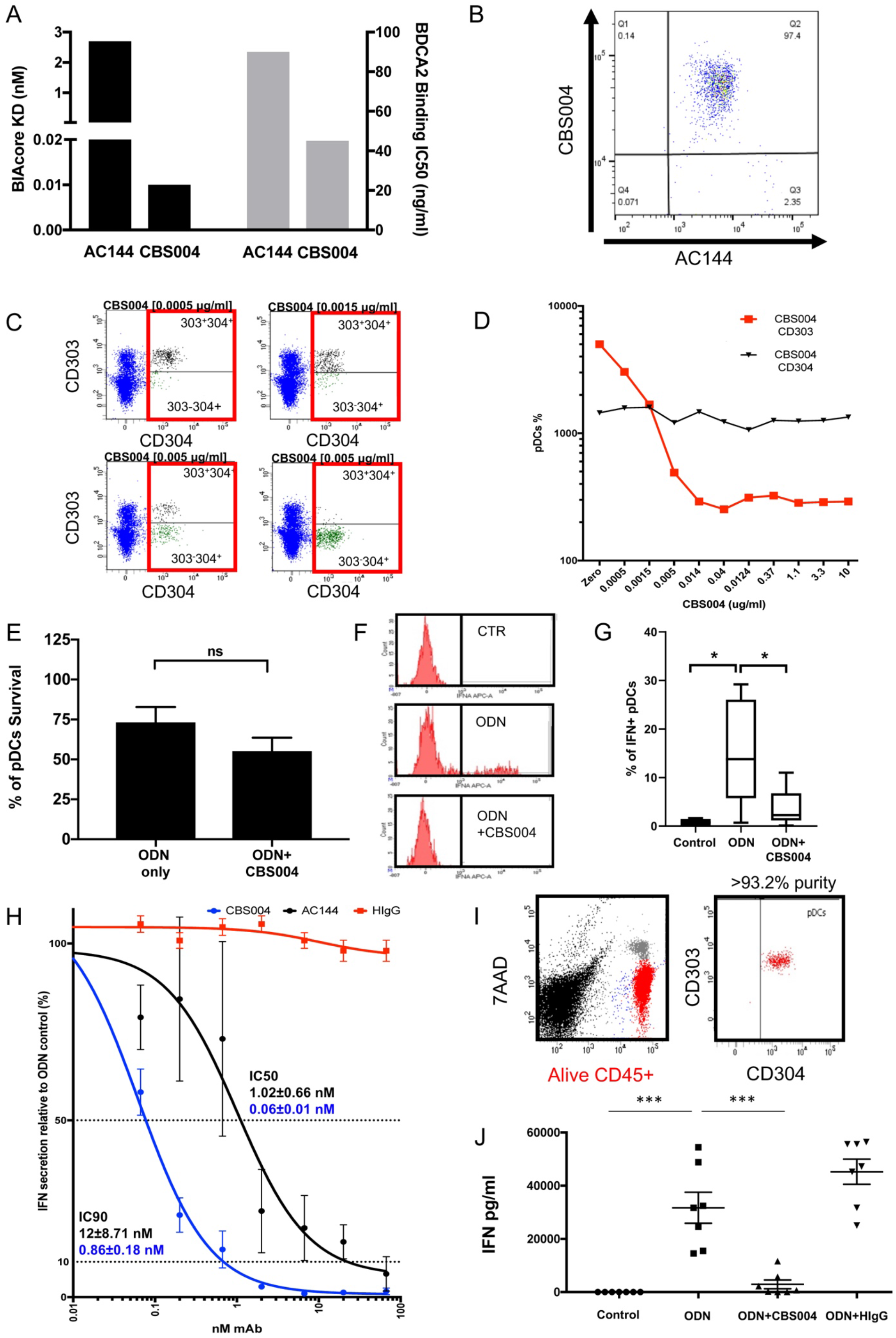
CBS004 specifically targets BDCA2 and blocks TLR9 induced human pDC IFN secretion. (A) Human BDCA2 binding affinity of CBS004 compared to miltenyi biotec AC144 antibody; BIAcore was used to determine the kinetic dissociation (KD, nM) of antibodies to BDCA2 (black bars); Competition binding ELISA showing the affinity of antibodies to BDCA2 (grey bars). Unbound BDCA-2-Fc protein at a dilution range [1-0.001 *μ*g/ml] was used to inhibit binding of CBS004 or AC144 [0.04 *μ*g/ml] to bound BDCA-2-Fc. IC50 was calculated by dose-response (grey bars). (B) CD304+CD123+ pDC gated according to Fig. S1 were stained with CBS004 and AC144 and analysed via FACS to illustrate alternate BDCA2 epitopes. (C) Illustrative CD303/BDCA2 internalisation FACS analysis of pDC gated from PBMC stained for CD303 and CD304 when cultured with increasing doses of CBS004 (from left to right; 0.0005, 0.0015, 0.005, and 0.014 *μ*g/ml). (D) Dose-response of increasing CBS004 concentration [0.005-10] on the mean fluorescence intensity (MFI) of pDC with surface bound CD303/BDCA2, normalised for CD304 MFI. Representative data from 3 HV. (E) Viability test of pDC within PBMC (n=11) cultured with ODN (1 *μ*M), and with CBS004 [10 *μ*g/ml] by measuring 7AAD staining via FACS. (F) Representative histogram of intracellular IFN alpha staining of pDC gated within PBMC when cultured in RPMI alone, with ODN (1 *μ*M), and with CBS004 [10 *μ*g/ml]. (G) Percentage of IFN positive pDC from FACS analysis from F (n=6). Percentage of IFNalpha secretion, measured by ELISA, from PBMC from 4 donors stimulated with ODN in the presence of CBS004, AC144 and HIgG [0-66 nM] relative to ODN-stimulated pDC with no antibody (100%). Dotted lines highlight IC50s and IC90s. Mean IC50 and IC90 values ±SEM displayed for CBS004 (blue) and AC144 (black). (I) Representative report of FACS analysis for purity of pDC isolated from HV PBMC using 7AAD-, CD45+, CD123+ gating strategy (Fig. S1). (J) IFN secretion from purified HV pDC (n=7) after 16 h of culturing in RPMI alone (CTR), with ODN (1 *μ*M), and with CBS004 or human IgG1 [10 *μ*g/ml] measured by ELISA. All results represent mean ±SEM. For (G) and (J) statistical significance was evaluated using unpaired two tailed t test. \**P* <0.05 and \*\*\**P* <0.001.

To determine whether CBS004 induced BDCA2 internalization we set out to measure the mean fluorescence intensity (MFI) of BDCA2 (CD303) as measured by bound AC144 on pDC gated within PBMC (Fig. S1) and normalizing for pDC cell number by using MFI of BDCA4 ab (CD304), another marker of pDC [23] (Fig. 1C). Treatment with CBS004 led to a dose-dependent decrease in BDCA2 surface expression on pDC, suggesting BDCA2 internalization (Fig. 1C and D). At 14 ng/ml of CBS004, internalization of BDCA2 reached saturation (Fig. 1D). Additionally, we showed that To BDCA2 targeting does not significantly reduce pDC viability as determined by 7AAD assay (Fig. 1E).

Functionally, ODN led to a 19-fold increase in IFN positive pDC (0.78±0.34% vs 14.98±4.40%) within PBMC (Fig. 1F-G, Fig. S1). CBS004 suppressed ODN-induced increase in IFN positive pDC by 96% (14.98±4.40% vs 3.73±1.62%), levels comparable to non-stimulated pDC (Fig. 1F-G).

To determine the ability of CBS004 to functionally suppress IFNα secretion of ODN-treated PBMC from healthy volunteers (HV) and compared to AC144 and human IgG controls (HIgG), we performed a dose-response in IFNα secretion measured in the cell supernatants by ELISA (Fig. 1H). CBS004 showed a 17-fold higher IFNα production inhibitory activity compared to AC144 (IC50 0.06 vs 1.02 nM, respectively) (Fig. 1H). When looking at the dose inducing 90% inhibition of IFN secretion, CBS004 induced a 14-fold more potent reduction of IFN compared to AC144 (IC90 0.86 vs 12 nM, respectively) (Fig. 1H). HIgG did not suppress IFN production.

To further validate these findings, we performed IFN secretion assays on pDC purified from PBMC, as previously described [19] which yielded >93.2% pure pDC (Fig. 1I). ODN stimulation induced a striking increase in IFNα secretion (31689±5809 vs 6.8±3.5 pg/ml, P=0.001), as expected (Fig. 1J). Incubation with CBS004 suppressed IFN secretion by 90% (2934±1639 pg/ml, P=0.0005), whereas [24]HIgG had no significant effect (Fig. 1J)

Jahn *et al.* have shown that BDCA2 internalization and inhibition of IFN secretion are dependent on receptor cross-linking with the Fc region of AC144 mAb, and monovalent binding of anti-BDCA2 Fab fragments was unable to inhibit ODN-induced IFN secretion, even at 50 nM [24]. To determine whether CBS004 inhibitory mechanism is dependent on Fc cross-linking, we generated digested Fab fragments of CBS004. Digestion of the fragments (50 kDa) was confirmed by SDS-PAGE gels using both non- and reducing loading buffers and visualized using Coomassie blue (Fig. S2A). Purified Fabs were quantified and used within the same experimental settings as in Fig. 1H. CBS004 Fab was able to reduce IFN secretion in a dose-dependent manner with IC50 of 18 nM (Fig. S2B). Nevertheless, the Fc-containing full IgG equivalents had a lower IC50 (Fig. S2B). This observation suggests that the epitope targeted by CBS004 is able to at least partially induce CD303 engagement, independently of the FC portion.

### CBS004 suppresses spontaneous and TLR-induced IFN secretion in SSc PBMC

Purified SSc pDC have been shown to produce higher levels of IFN compared to HV [10, 11]. Consistent with these findings, we found that PBMC from SSc patients showed an average basal level secretion of IFNα 25.13±12.38 pg/ml compared to undetectable levels from HV PBMC (Fig. 2A). ODN stimulation of PBMC induced a substantial increase in IFN secretion in both HV and SSc PBMC (49410±17359 pg/ml and 67494±27958 pg/ml P<0.05, respectively) (Fig. 2A). CBS004 treatment suppressed the ODN-induced IFN secretion by >98% in all samples (HV 555.1±355 pg/ml and SSc 1324±614.8 pg/ml, P<0.05, respectively) (Fig. 2A).

**Figure 2.**
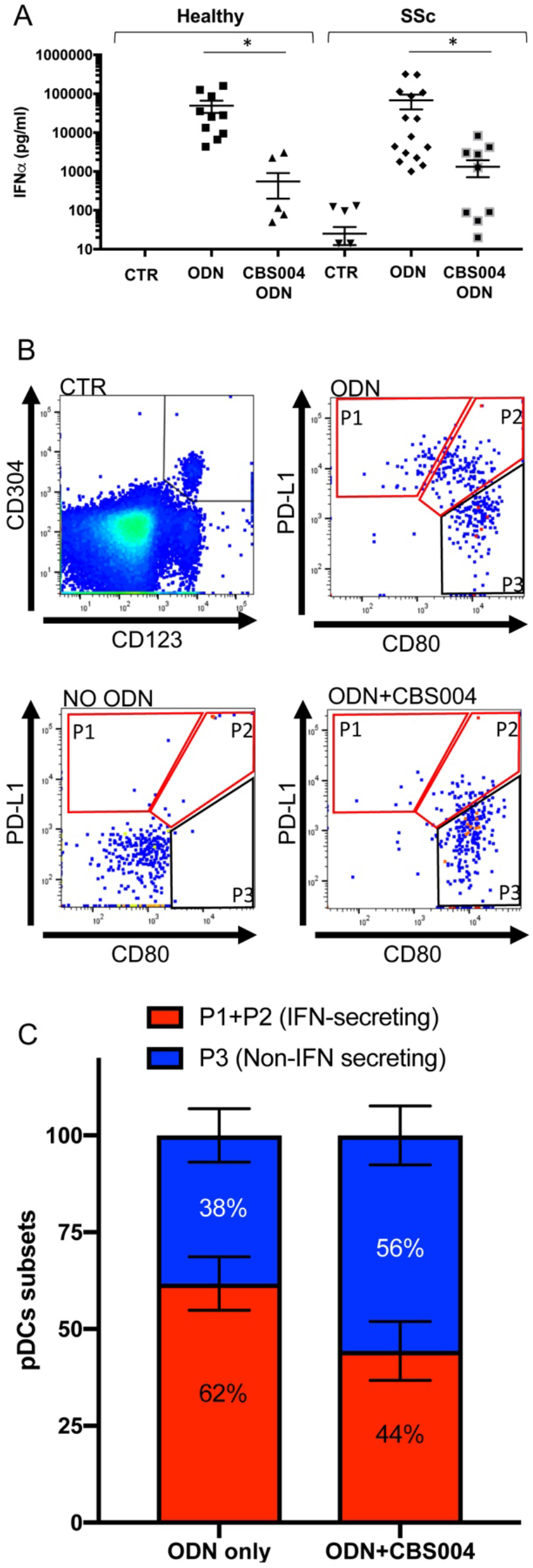
CBS004 suppresses spontaneous and TLR-induced IFN secretion in SSc PBMC. (A) PBMC purified from healthy and SSc donors were cultured in media alone (CTR), with 1 *μ*M ODN, or with ODN and CBS004 [10 *μ*g/ml] for 16 h (n=15). IFNalpha was quantified in the supernatants by ELISA. (B) Example of sub-typing FACS analysis of pDC (CD123^+^CD304^+^) in the 3 culture conditions (CTR, ODN, ODN+CBS004). Gating illustrates P1, P2, P3 subtypes. Red highlighted gates display IFN-secreting sub-types as previously described [25]. SSc PBMC were cultured for 16 h as in and pDC were sorted as Lin^−^HLA-DR^+^CD45^+^CD123^+^CD304^+^ (Fig. S1) and gated for PD-L1 and CD80 expression to determine P1, P2 and P3 sub-types. P1, PD-L1^+^CD80^−^; P2, PD-L1^+^CD80^+^; P3, PD-L1^−^ CD80^+^. [25]. (C) Quantification of sub-types based on FACS analysis of PBMC samples (n=7) between ODN and ODN+CBS004 culture conditions. All resulted are represented as means ±SEM. For (A) statistical significance was evaluated using unpaired two-tailed t-test. \**P* <0.05.

It has been recently shown that TLR-induced activation of pDC triggers stable cell differentiation into three stable sub-types, with PD-L1(CD274)^+^CD80^−^ (P1) and PD-L1(CD274)^+^CD80^+^ (P2) subpopulations specialized in type I IFN production both in healthy controls and patients with autoimmune conditions [25]. To determine the functional consequences of BDCA2 targeting ODN-activated pDC in SSc, we measured the proportion of PD-L1 and CD80 positive subpopulations (Fig. 2B-C). Similarly, to what has been observed following viral stimulation [25], treatment with ODN induced pDC to differentiate in the specific subtypes, with 62% of pDC differentiating in P1 and P2 (Fig. 2B-C). In this context, treatment with CBS004 showed a 1.4-fold reduction in the IFN-producing pDC subsets (Fig. 2B-C).

### CBS004 driven targeting of BDCA2 suppresses overall pDC transcriptome activation

Transcriptome profiling of immune cell populations, including DC subsets, has provided unprecedented new insights into pDC biology. To determine the overall effects of BDCA2 targeting on pDC transcriptome, we performed RNA-seq analysis of four independent donors of human pDC following ODN-induced TLR-9 stimulation with or without BDCA2 targeting via CBS004. Transcriptome analysis revealed 328 Differentially Expressed Genes (DEGs ≥ or ≤2-fold change; FDR ≤ 0.05) between unstimulated and ODN-stimulated pDC (Fig. 3A and STab.1). Pathway analysis on this set of DEGs identified genes involved in ‘response to virus’, defence response to other organism’, and ‘defence response to virus’ at the top of the enriched biological processes (Fig. 3B).

**Figure 3.**
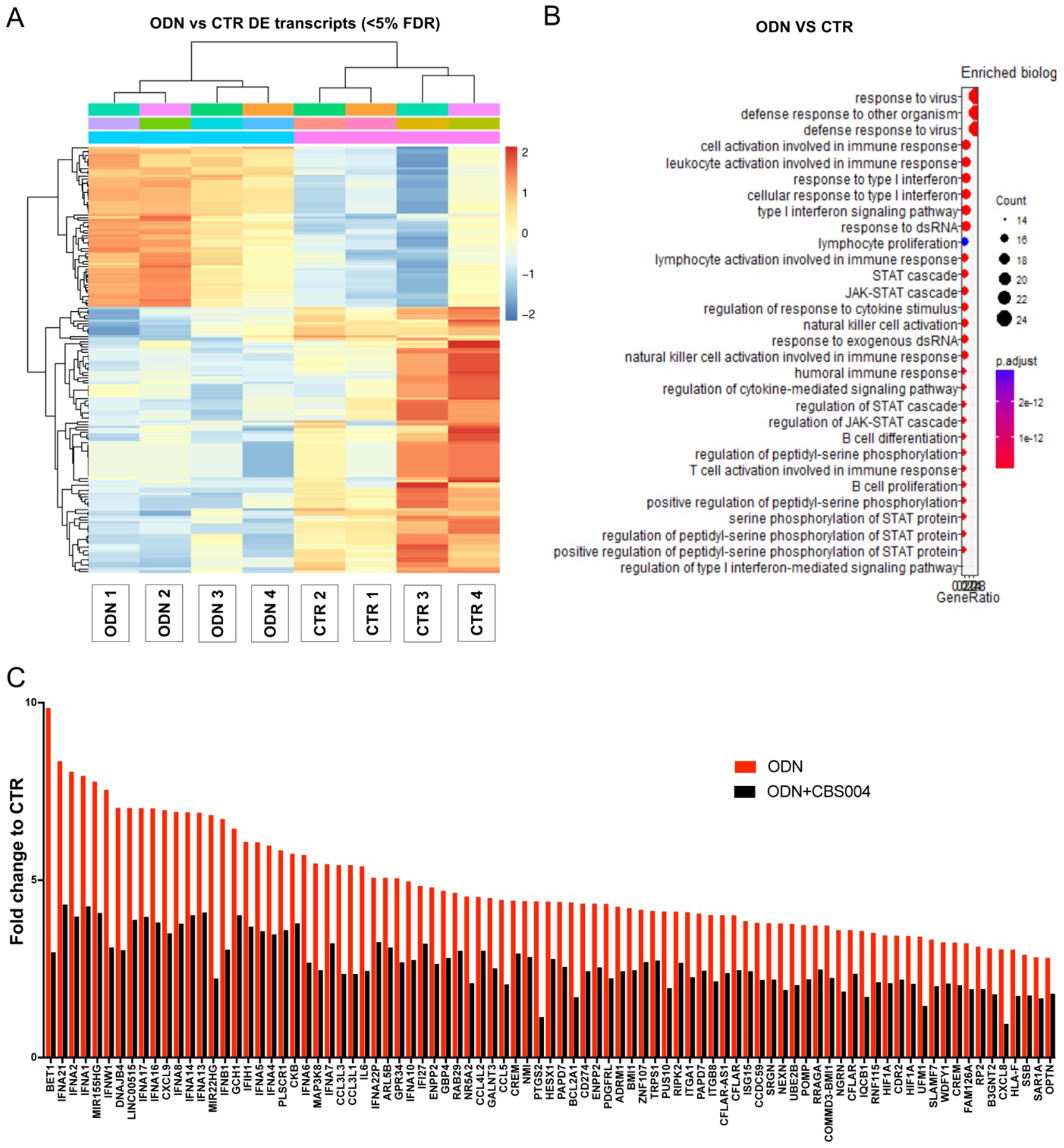
CBS004 driven targeting of BDCA2 suppresses overall pDC transcriptome activation. Transcriptome analysis of human pDC cultured in media alone (CTR), with 1 *μ*M ODN2216 (ODN), or with ODN and CBS004 [10 *μ*g/ml] for 16 h (n=4). (A) Heatmap of reduced, centered normalised read counts for differentially expressed (DE) transcripts among CTR and ODN populations, <5% False Discovery Rate (FDR), calculated using Benjamini-Hochberg multiple testing correction. DE transcripts ≥ or ≤2-fold (FDR <0.05) shown in STab.1. (B) KEGG pathway analysis showing top biological processes enriched in the set of DE transcripts between CTR and ODN pDC, and their associated P values. (C) Average fold change of DEGs for ODN and ODN+CBS004 relative to CTR (n=4). For repeat transcripts, highest fold change used for comparison. Red bars represent the 87 genes that were increased ≥2-fold (FDR <0.05) between CTR and ODN that are dependent on BDCA2 treatment (reduced ≥1.5 fold by CBS004) (full transcript data, STab.2).

The biological processes shown to be ODN-inducible within pDC match to those previously identified in characterised inflammatory SSc skin subsets, suggesting that pDC activation may be involved in this specific inflammatory subset. Consistent with this notion, we observed upregulation of many IFN-related genes, such as IFNA2, IFNA21, IFNB1, and CCL5, a common feature seen in SSc. Pathway analysis also showed JAK/STAT, NF-kB and angiogenesis pathways to be major biological processes upregulated by ODN-stimulation (Fig. 3B), which have been shown to be dysregulated in SSc, but not shown in pDC before. These data suggest that TLR-stimulation of pDC can induce a multitude of genes beyond IFN, which could contribute to the pathogenesis of tissue damage in SSc.

ODN treatment of pDC induced the expression of 145 genes ≥2-fold (FDR ≤0.05) (STab.1). Pre-treatment with CBS004 reduced the expression of 60% of ODN-inducible DEGs ≥1.5 fold (Fig. 3C and STab.2). CD274 expression was ODN-induced and dependent on BDCA2 targeting, which supported ours and others’ observations that 12-18 h TLR stimulation induces P1 and P2 subsets that are CD274^+^ and secrete IFN [25]. ODN-induced IL6 expression was also dependent on BDCA2 targeting. Interestingly, IL6 and IFN secretion can synergistically activate B cells [25, 32]. Furthermore, one of the ODN-inducible and BDCA2-targeted genes was proteoglycan serglycin (SRGN), which has been shown to be secreted into the extracellular matrix (ECM), and linked to promoting lymphoid cells adhesion and activation [33, 34], storage of chemokines and cytokines, as well as being able to induce epithelial-mesenchymal transition (EMT) [35,36]. Together, these data demonstrate that TLR-stimulation of human pDC goes beyond IFN secretion induction and predicts a greater biological relevance of pDC activation in SSc pathogenesis. More importantly, our analyses show that TLR-induced pDC transcriptome can be drastically suppressed by targeting of BDCA2.

### BDCA2 targeting with CBS004 suppresses the pDC induced IFN signature of organotypic skin grafts

Growing evidence shows SSc patients have pDC skin infiltration and induced IFN signature within the skin 10–12]. To determine the IFN-induced skin response to human pDC we exploited organotypic 3D skin rafts (OSRs) as an *in vitro* model that mimics the microenvironment of the skin and allows cross-talk between the two main cellular components, fibroblasts and keratinocytes (Fig. 4A). OSRs were treated with supernatants from pDC cultured in RPMI alone (CTR), pDC stimulated by ODN (ODN), or ODN+CBS004 (ODN+AB). To measure the pDC-dependent IFN-induced transcriptome we interrogated by qRT-PCR the gene expression levels of 78 key interferon signalling genes (ISGs) in three independent OSRs preparations. ODN-conditioned media resulted in 35 ISGs upregulated between 1.8-32-fold in the OSRs, relative to expression in CTR OSRs (Fig. 4B). Interestingly, we observed a high inter-donor variability as far as magnitude of gene-induction between triplicate experiments. The 8 genes more consistently upregulated included *ISG15*, *IFITM1*, *BST2*, *IFI6*, *IFIH1*, *NMI*, *HLA-B* and *IFITM3* (3-19 fold induction relative to control and P<0.05), showing that TLR-activated pDC alone can stimulate an IFN response within the skin. Most importantly, conditioned media from ODN+AB resulted in suppressed upregulation in all of those genes with a reduction in their expression ranging from 1.8 to 11-fold compared to gene expression induced by ODN (Fig. 4C). This resulted in a transcription profile similar to the control condition (CTR OSRs - Fig. 4D). Together these results suggest that specific BDCA2 targeting of pDC can suppress the induced IFN response of skin cells, providing mechanistic evidence that pDC inhibition could reduce the interferon response seen in SSc.

**Figure 4.**
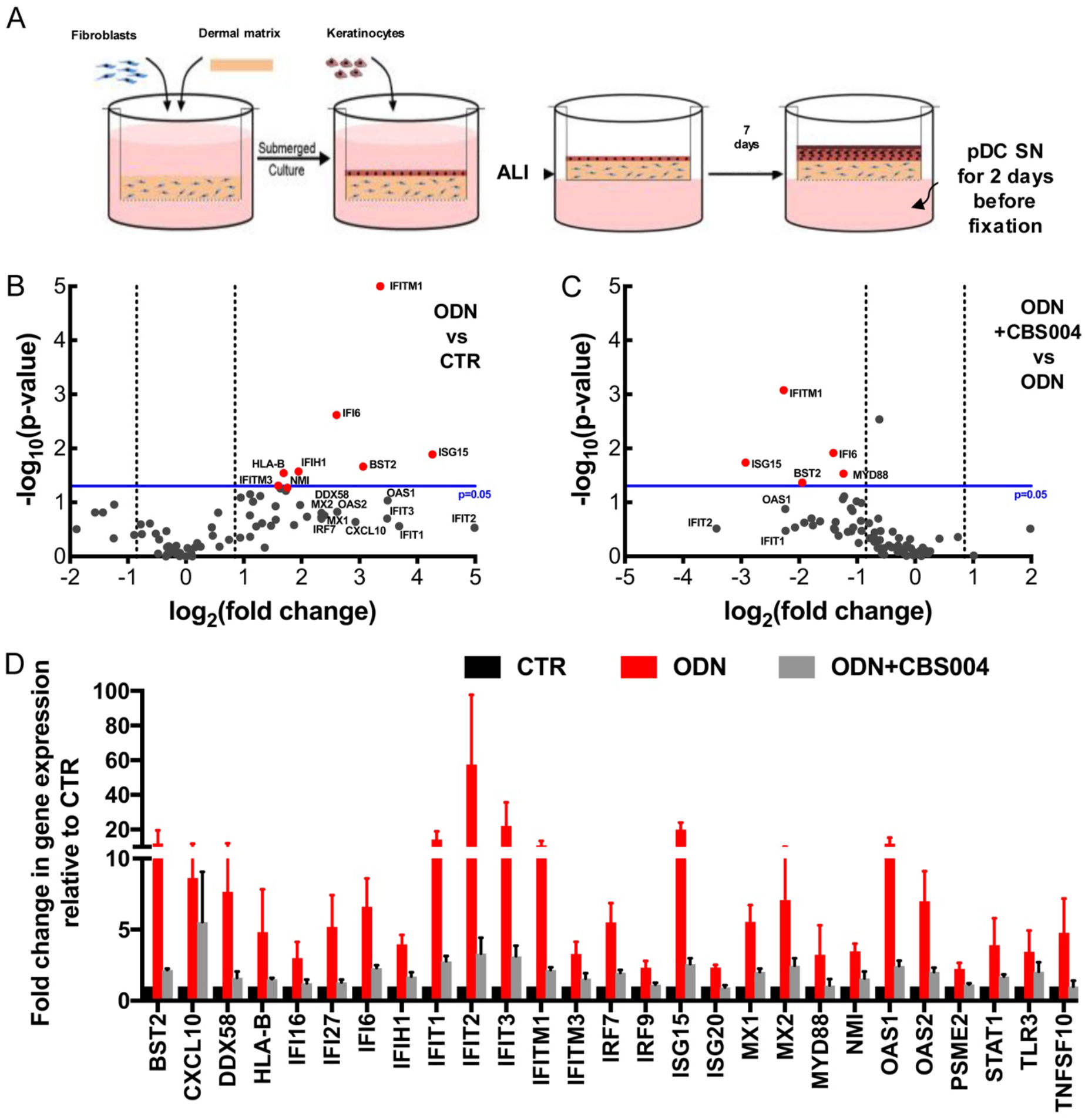
BDCA2 targeting with CBS004 suppresses the pDC induced IFN signature of organotypic skin grafts (OSRs). (A) Systematic outline of OSR protocol; fibroblasts are embedded into a collagen matrix and keratinocytes seeded above until confluence. OSR is brought to air-liquid interface (ALI) to sustain epithelium differentiation. After 5 days, ALI media spiked with 6000 pg/ml of IFN (generated by TLR-stimulated pDC; ODN) for 48 h. CTR; contains equivalent supernatant from untreated pDC (undectable IFN) and from pDC treated with ODN+CBS004 [10 *μ*g/ml]. (B and C) RNAs harvested from 3 mm biopsies from OSRs and subjected to type I IFN inducible genes superarray. Volcano plots illustrate the fold change of 79 IFN type I-related genes (black dots) between CTR and ODN (B) and between ODN and ODN+CBS004 (C) (n=3 for each condition). Grey lines represent the 1.8-fold change cut off. Blueline represents the cut off for statistical significance of *P* 0.05 calculated using a Student’s t-test (two-tailed distribution and equal variances between the two samples) on the triplicate 2-∆Ct values for each gene in each treatment group compared to the control group. (D) Bar chart illustrates the IFN type I-related genes that were > 1.8 fold increased in ODN relative CTR and the effect of CBS004. Results are represented as means ± SEM.

### Xenotransplant of human pDC in NOD Scid mice increased skin IFN response to TLR-stimulation in a BDCA2-dependent manner

To determine whether pDC were responsible for eliciting an ISG response in the skin microenvironment *in vivo* following exposure to disease-relevant activators of inflammation, we developed a novel xenotransplant model. This model involved transfer of purified normal human primary pDC into immune-deficient NOD Scid mice (XenoSCID) via intravenous (i.v.) injection followed by topical application of imiquimod-containing cream (Aldara), with or without anti-BDCA2 (CBS004) or human IgG (HIgG) (Fig. S3). The purity of pDC isolated from healthy PBMC and functional responses to TLR-9 were assessed for each batch (Fig. 1I and S4). Skin samples were taken at 12 hours post i.v. pDC injection, and enzymatically digested to release cells. FACs analysis of CD45^+^CD123^+^CD304^+^cells indicated pDC skin infiltration within the Aldara-treated skin (0.3% of total cells, Fig. 5A). Most importantly infiltration of skin with human pDC resulted in a functional increase in ISG expression as assessed by qRT-PCR analysis of 78 genes commonly upregulated during a type I Interferon response. Mice receiving human pDC had at least 2-fold upregulation of 15 mouse type-I IFN response genes including *Ccl5*, *Cxcl10*, *Ifit1*, *2* and *3*, *Isg15*, *Mx1* and *2*, *Oas1a* and *1b*, and *Stat1* and *2*, compared to Aldara treatment alone in absence of pDC (P<0.004) (Fig. 5B).

**Figure 5.**
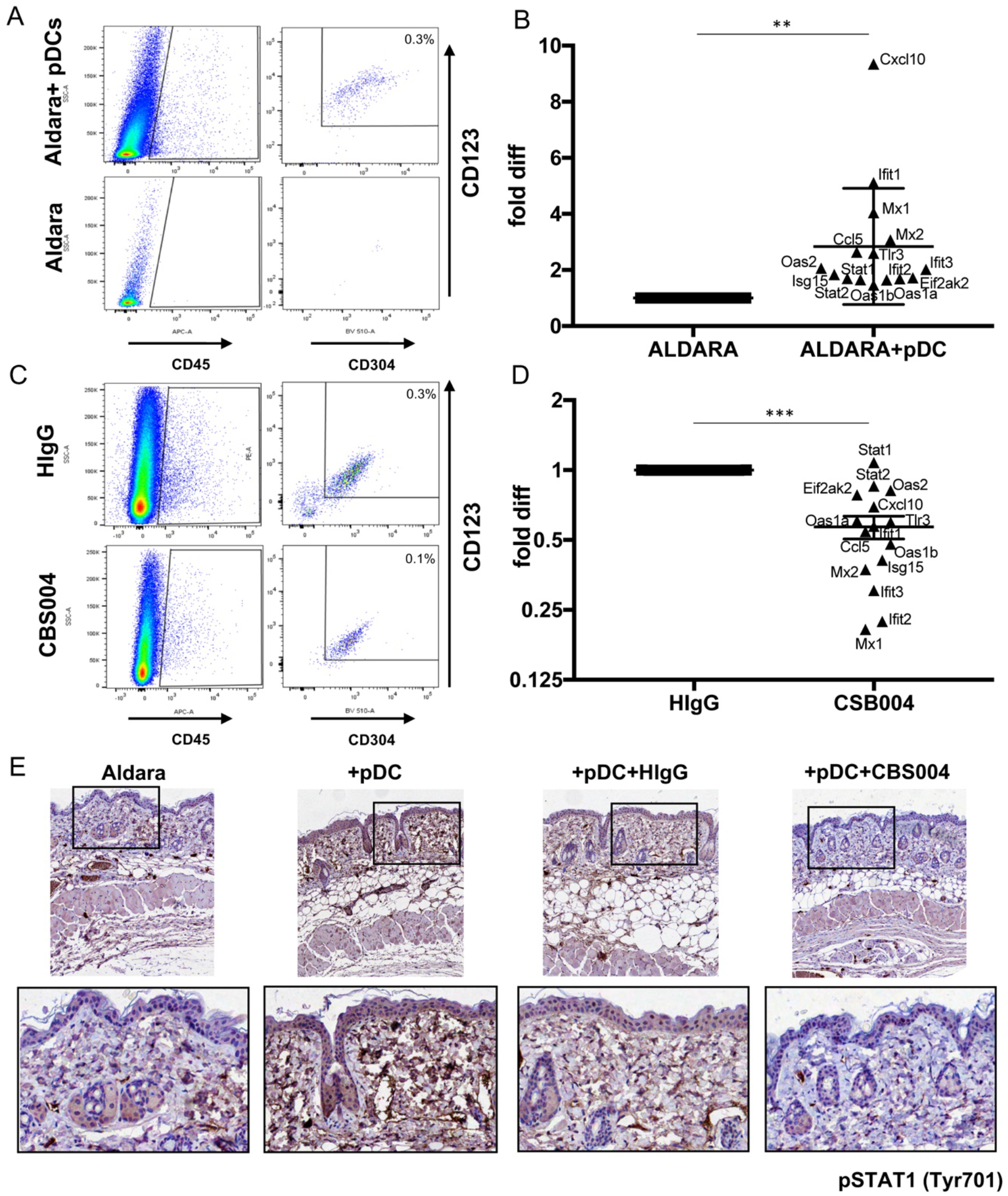
XenoSCID with human pDC increased skin IFN response to TLR stimulation in a BDCA2-dependent manner. Intravenous (i.v.) tail injection of 2.5×10^5^ human purified pDC and intraperitoneal (i.p.) injection of CBS004 mAb (5 mg/kg) or control human IgG to NOD Scid mice treated with topical Aldara cream application (Fig. S3; systematic diagram and timeline), with four different treatment conditions consisting of Aldara, Aldara+pDC, Aldara+pDC+CBS004 and Aldara+pDC+HIgG, each in duplicates. Treated skin was harvested using a 3 mm punch biopsy and processed for FACS analysis of human pDC (CD45^+^CD123^+^CD304^+^) (representative analyses A and C), qRT-PCR analysis for type I IFN inducible genes (B and D), and IHC staining for pSTAT1 (Tyr701) (representative images E). (B) Mouse specific type I IFN inducible genes increased >2-fold within the skin of mice between Aldara and Aldara+pDC conditions, represented as a global mean ±SEM with individual genes highlighted. (D) Illustrates the changes in expression of pDC-inducible genes from (B). For (B) and (D) statistical significance was evaluated using paired two-tailed t-test. \*\**P* <0.01 \*\*\**P* <0.001.

In this context we could assess the *in vivo* efficacy of BDCA2-targeting. CBS004 and HIgG control mAb were injected into XenoSCID 12h before pDC i.v. Injection and the ability of the mAbs to inhibit human Aldara-stimulated pDC examined. Aldara-induced pDC skin infiltration, as detected by alive human CD45^+^CD123^+^CD304^+^ cells in the mouse treated skin, was not reduced by HIgG (0.3% of total cells), however was reduced by 3-fold to 0.1% with CBS004 treatment (Fig. 5C). Most importantly, XenoSCID receiving CBS004 suppressed 93% of the pDC-inducible type-I IFN response genes compared to mice treated with HIgG (Fig. 5D). Consistent with these findings, IHC analysis for pSTAT1 (Tyr701) protein levels showed a strong induction by pDC i.v., which was dramatically reduced by BDCA2-targeting and unaffected by HIgG administration (Fig. 5E). Together the above data demonstrate that XenoSCID is a viable animal model to investigate human pDC induced skin IFN response to TLR agonists and that BDCA2-targeting can suppress the pDC induced skin IFN response in this model.

### Xenotransplant of human pDC in NOD Scid mice increased the skin profibrotic response to bleomycin treatment in a BDCA2-dependent manner

Ah Kioon *et al.* have shown that depletion of mouse pDC can ameliorate bleomycin-induced skin fibrosis. To determine the role of the human specific BDCA2 in this setting we developed our XenoSCID model with bleomycin-induced skin fibrosis. In this setting, we supplemented every other day s.c. injection of bleomycin with weekly tail vein injection of human pDC for 3 weeks (Fig. S5). As anticipated, bleomycin alone induced a blunted fibrotic response at three weeks, as shown by partially retained fatty layer and no significant increase in overall skin thickness when measured using haematoxylin and eosin staining (Fig. 6A). Accordingly, Masson trichrome staining (Fig. 6B) and epidermis and dermal thickness (Fig. 6C) were only mildly affected. Consistent with these observations, SIRCOL analysis of collagen content within skin samples also showed no significant difference between control and bleomycin alone treatments (Fig. 6D). On the contrary, the presence of human pDC resulted in the bleomycin-induced loss of all subdermal fat, along with increased collagen formation (measured by SIRCOL) and a 40% increase in overall skin thickness (Fig. 6A-D). The fibrotic response was associated with Type I IFN signalling activation as suggested by increased MX1 and pSTAT1 (Tyr701) protein expression with i.v. of pDC (Fig. 6E and F).

**Figure 6.**
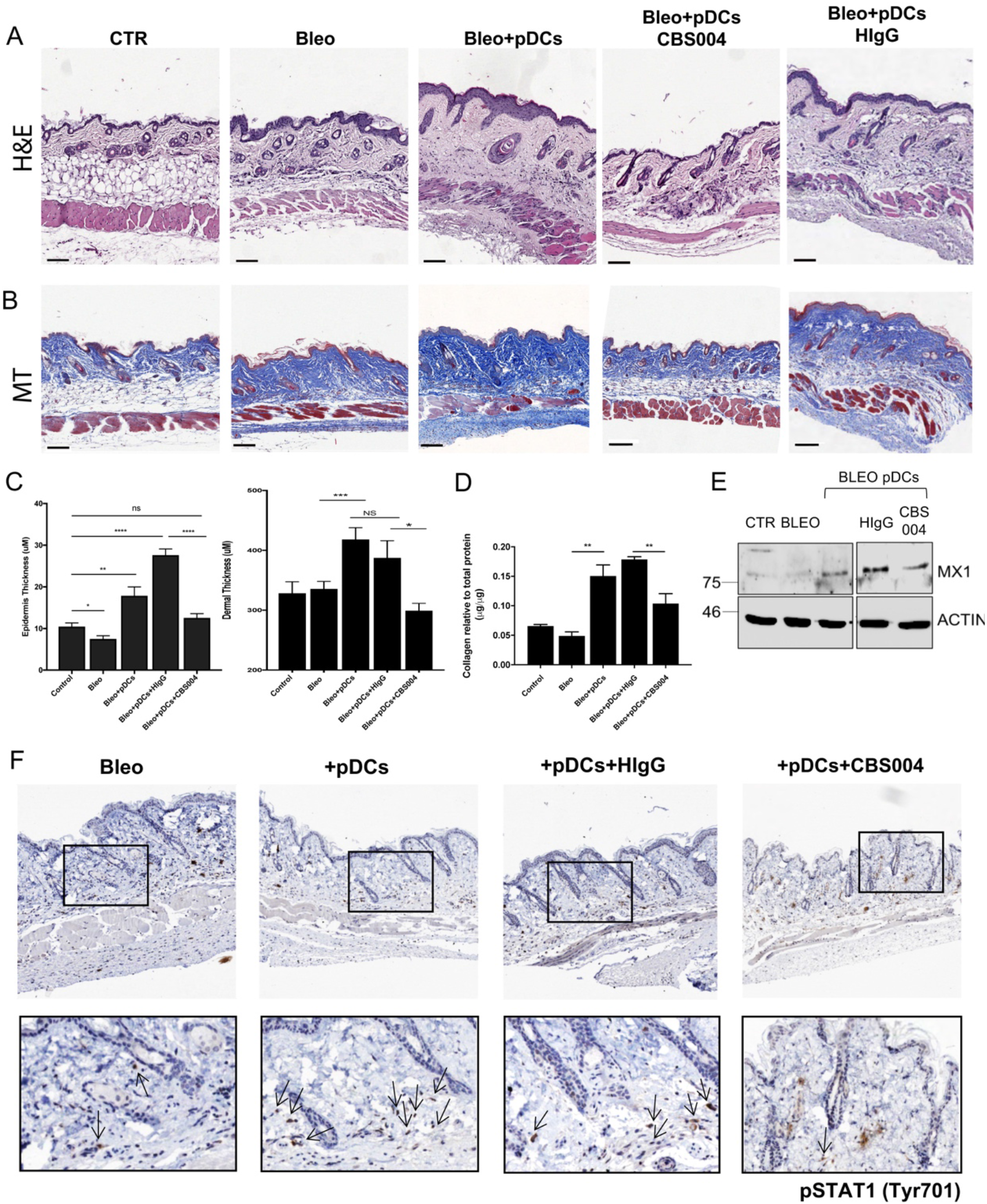
XenoSCID with human pDC increased the skin profibrotic response to bleomycin treatment in a BDCA2-dependent manner. Intravenous (i.v.) tail injection of 2.5×10^5^ human purified pDC and intraperitoneal (i.p.) injection of CBS004 mAb (5 mg/kg) or control human IgG into NOD Scid mice treated with bleomycin (Bleo) or PBS injections (Fig. S5; systematic diagram and timeline), with five different treatment conditions consisting of PBS/Control, Bleo, Bleo+pDC, Bleo+pDC+CBS004 and Bleo+pDC+HIgG, each in triplicates. Treated skin was harvested using a 3 mm punch biopsy and processed for Hematoxylin and Eosin (H&E) (A) and Masson Trichrome (MT) staining (B). (C) Epidermis and dermal thickness were measured from 20 areas in each condition. (D) An additional punch biopsy was taken and used to extract protein. Total collagen content measured by SIRCOL assay and shown relative to total protein concentration. Results represented as means ± SEM of triplicate experiments. Statistical significance was evaluated using paired two-tailed t-test. \**P* <0.05 \*\**P* <0.01 \*\*\**P* <0.001 NS=no significance. (E) Western blot analysis of MX1 and BETA-ACTIN protein expression. (F) IHC analysis of pSTAT1 (Tyr701) representative images with zoomed in areas, arrows highlight positively stained cells.

To determine the therapeutic implications of BDCA2 targeting in this setting, bleomycin stimulated XenoSCID were treated with i.p injection of CBS004 or Human IgG (Fig. S5). pDC-induced skin fibrosis was dramatically reduced by administration of CBS004 as demonstrated by the retention of some fatty layer tissue, similar to bleomycin-only treated mice, a 2-fold reduction in dermal and epidermal thickness and 1.5-fold reduction in collagen content compared to HIgG (Fig. 6A-D). Furthermore, the therapeutic effects of BDCA2-targeting were associated with a reduction in MX1 compared to HIgG treatment (Fig. 6E). The same was true with specific pSTAT1 dermal fibroblast (based on morphology) protein expression (Fig. 6F).

Overall our Xenotransplant models using human pDC clearly show that TLR-activated pDC have a crucial role in skin inflammation and fibrosis. By using BDCA2-specific pDC targeting, these major phenotypes relevant to SSc were greatly repressed supporting this approach as a viable therapeutic option for the disease.

## Discussion

Elevated IFN gene signature in affected organs and in the blood is a key indicator of SSc disease severity, and has been shown to precede the onset of clinical fibrosis [13]. Since pDC are known to be the key cell type for activating IFN response to virus [3, 4], along with increasing evidence suggesting mouse pDC are important for disease development [10, 37], we developed our study to research the role of human pDC in inflammation and fibrosis of the skin. Specifically, we set out to determine whether targeting BDCA2 could suppress the IFN response and pro-fibrotic activation of pDC and ultimately inform the rationale for pDC targeted therapy for SSc.

Our study clearly shows that human pDC can be TLR-activated to secrete IFN and to a gene expression profile mapping to activation of inflammation, JAK/STAT, NF-kB and angiogenesis pathways, predicting a greater biological relevance of pDC activation in SSc pathogenesis. Furthermore, we have shown that BDCA2-targeting using CBS004 mAB is effective at blocking *ex vivo* pDC IFN secretion and ODN-induced transcriptome.

By using organotypic models of human skin we were able to show that TLR-activated human pDC could stimulate a skin IFN response, in a BDCA2 dependent manner. Our xenotransplant model using human pDC greatly expanded this observation. By inducing local TLR-activated skin, we have shown that human pDC can infiltrate this site and induce a mouse IFN response within skin. Furthermore, we clearly see that human pDC are capable of inducing IFN and fibrotic skin response when introduced into our bleomycin mouse model. Together, our data indicate that human pDC, and their IFN production, can be a key cell type in the pathogenesis of skin IFN response and fibrosis in SSc. BDAC2 targeting of pDC in situ prevented the pathogenic responses to IFN and pro-fibrotic stimuli, identifying specific pDC targeting to be a viable therapeutic application for SSc.

Previous reports showed that that BDCA2 targeting pDC inhibition was dependent on both FAB and FC portion of the antibody [24]. Nevertheless, here we have highlighted that the epitope targeted by CBS004 has a more potent inhibitory effect, since the FAB-BDCA2 engagement also inhibited IFN production by activated pDC, although with reduced efficacy.

Since the first reports on the inhibitory effects of BDCA2 targeting, it has now been proposed that pDC differentiate in different functional classes following activation [25]. Here we show that BDCA2 engagement strongly suppresses the differentiation of IFN-secreting CD274+ pDC with a prevalent differentiation towards CD274^−^CD80^+^ pDC. Further functional studies will shed light on the effects of BDCA2 engagement on T cell co-stimulation, which has been suggested to be affected by CD274^−^CD80^+^ pDC [25]. Our RNA-seq data also expand on this notion showing that the inhibitory effect of BDCA2 goes beyond IFN production and extends to the majority of the transcriptome induced by TLR-9 engagement.

The development of our XenoScid model is a novel tool that can be used to study the biology of human pDC in mice and can be applied in the research of other IMIDs affecting the skin, such as Psoriasis or Cutaneous Lupus Erythematosus (CLE). A limitation of this approach is the lack of adaptive immune response in these animals. Therefore, the consequence of pDC inhibition in a competent immune system remains unknown. Nevertheless, the studies from Rowland *et al.* showed that in a mouse model of lupus, elimination of pDC strongly impaired expansion and activation of T and B cells [38]. In this context, xenotransplant models of human PBMC with and without pDC-depletion would be extremely informative, although falling beyond the scope of this study.

In conclusion, we have presented data that support the role of human pDC in inducing an inflammatory and fibrotic response within the skin, further implicating pDC in SSc pathogenesis. Most importantly we show that the blocking of pDC function by BDCA2-targeting prevented pathogenic responses to IFN and pro-fibrotic stimuli, as shown in cultured cells, skin rafts and *in vivo*, supporting the development of this approach as therapeutic application for SSc and other IMIDs.

## Material and Methods

### Study design

The research objective of our study was to determine whether BDCA2 targeting is sufficient to inhibit TLR induced pDC activation. We developed a xenotransplant model of human pDC in immunocompromised mice (XenoSCID) and defined the effects of pDC and BDCA2 targeting in TLR-induced skin inflammation and bleomycin-induced skin fibrosis to understand whether targeting pDC could be a potential therapeutic opportunity for SSc and other pDC-implicated autoimmune diseases.

### Affinity analysis of humanized mAbs by BIAcore

A BIAcore T200 was used with BIAcore run buffer (HBS-EP) at pH7.4. 692RU of huBDCA2-Fc was immobilized to a CM5 chip (CFJB156) utilising 5 *μ*g/ml of huBDCA2-Fc with the BIAcore EDC/NHS kit according to the manufacturer’s instructions. Two-fold dilutions of the anti-human CD303 BDCA2 antibody (AC144) (Miltenyi Biotec), chimeric CBS004 (Capella Bioscience) or humanized CBS004 mAbs (LONZA, Cambridge) were injected starting at 200 nM down to 3.1 nM with a contact time of 60s at a flow of 30 or 60ul/min at 25C followed by an off-rate wash for 5 minutes with BIAcore buffer. Regeneration of the chip was achieved with two injections of 10*μ*l of 10 mM NaOH/1M NaCl between samples. The BIAcore T200 software was used to calculate Ka (1/Ms), Kd (1/s) and the KD (nM).

### Competition ELISA assay

Wells were coated with 0.5 *μ*g/ml of human (hu) BDCA2-Fc in 1x PBS pH 7.4 O/N at 4°C, added at 100 *μ*l per well. Blocking was achieved with 4 % Skimmed Milk (Marvel) in PBS for 2 h at RT. To the well, 50 *μ*l of huBDCA-2-Fc protein was added in a dilution range [1-0.001 *μ*g/ml] in 1 % Skimmed Milk in 1x PBS (buffer) plus 50 *μ*l of CBS004, or AC144 in buffer at [0.04 *μ*g/ml] was added for 1 h at RT. Bound CBS004 and AC144 were detected by secondary antibodies: Mouse Anti-Human IgG (anti-CH1-HRP) (BD Pharmigen) at 1:1000 or Peroxidase conjugated Affinipure Donkey anti-mouse IgG (anti-mouse IgG-HRP) at 1:5000 (Jackson ImmunoResearch), respectively, in buffer. Control 2nd antibody: anti-CH1-HRP and anti-mouse IgG-HRP in 1 % Skimmed Milk / 1x PBS; 100 *μ*l/well. Development achieved by TMB (ThermoFisher) and Stop reaction by H_2_SO_4_ and the absorbance read at 450nm.

### Peripheral blood mononuclear cells handling

Blood samples from 15 SSc patients enrolled at the Scleroderma clinic within the Leeds Institute of Rheumatic and Musculoskeletal Medicine (UK) and from 15 healthy subjects, matched for gender and age, were analyzed in this study. All SSc patients enrolled fulfilled the 2013 EULAR/ACR classification criteria for SSc. All subjects signed informed consent to participate in the study and all related procedures. Peripheral blood mononuclear cells (PBMC) were isolated from EDTA anti-coagulated blood by density gradient separation using prefilled Leucosep™tubes (Greiner Bio-One Ltd, UK). PBMC were cultured in RPMI1640 containing 10% FBS and 1% Penicillin Streptomycin (PS) (Gibco Laboratories, Grand Island, NY).

### FACS analysis of pDC

PBMC were labelled for FACS staining with mouse anti-human antibodies directed against lineage markers (Vioblue-CD3, CD14, CD19, CD56 and CD11c) (BD Biosciences), APC-Vio770 HLA-DR, PerCPVio770-CD123 (IL-3R) and PE-CD304 (BDCA4) FITC-CD303 (BDCA2) (Miltenyi Biotec). The plate was then incubated at 4 C for 30 minutes followed by addition of 200ul of FACS buffer and centrifugation at 300g for 10 minutes at 4 C. Supernatants were decanted, and cell wash repeated using 200ul of ice-cold Dulbecco’s PBS. Cell pellets were re-suspended in 200ul of Fixation/permeabilization buffer (e-Biosciences) and the plate incubated at 4 C for 30 minutes. Following centrifugation, cells were washed once in 200 ul of perm/wash buffer (e-Biosciences). Finally, the cells were re-suspended in FACS buffer for flow cytometry analysis. The data acquisition was performed on LSRII 4 laser flow cytometer (BD Biosciences), and the analysis was conducted using FACS DIVA software (BD Biosciences). pDC gating strategy excluded dead cells using Aminoactinomycin D (7-AAD) (BD Biosciences) and lineage-, and HLA-DR+, with sequential gating for human CD45+CD123+CD304+ (Fig. S1). For competition assay between CBS004 and AC144, both antibodies were incubated with fixed PBMC on ice for 30 minutes. For intracellular detection of IFN, prior to resuspending in FACS buffer, cells were labelled with APC IFN-I (Miltenyi Biotec) at 4◻C 30 minutes.

### BDCA2 internalisation experiments

PBMC (2*10^6 cells) were maintained for 16h in RPMI1640+10% FBS+ 1% PS in 96 well round bottom plate. Cells were cultured with or without 0.5 *μ*M for ODN2216 (Miltenyi Biotec) in presence of increasing concentration of CBS004 (0.0005-10 *μ*g/ml). The plate was centrifuged at 300g for 10 minutes and cells labelled for FACS staining with mouse anti-human antibodies directed against lineage markers, APC-Vio770 HLA-DR, PerCPVio770-CD123 (IL-3R) and PE-CD304 and FITC-CD303. The analysis gating strategy was aiming to detect change in mean fluorescence intensity of cell surface CD303 expression (detected with Miltenyi Biotec clone AC144) on pDC (CD3-18-56-14-CD11C-DR+CD123+CD304+).

### Isolation of plasmacytoid dendritic cells (pDC)

For healthy pDC, commercial cryo-human pDC (Stemcell Technology) from 5 healthy subjects, matched for gender and age, as well as pDC isolated from PBMC isolated from healthy cone donations, using Diamond Plasmacytoid Dendritic Cell Isolation Kit II (Miltenyi Biotec) following manufacturer protocol, were used. Firstly, the non-pDC are indirectly magnetically labeled with a cocktail of biotin-conjugated antibodies against lineage-specific antigens and Anti-Biotin MicroBeads. Followed by depletion of non-pDC, using an LD MACS® column and magnetic field MACS Separator (Miltenyi Biotec), the pre-enriched pDC were labeled with pDC-specific CD304 (BDCA-4/Neuropilin-1) Diamond MicroBeads and isolated by positive selection over a MS MACS Column and magnetic field MACS Separator (Miltenyi Biotec). Purified pDC were then counted and tested for purity by FACS staining with mouse anti-human antibodies directed against lineage markers (VioBlue-CD3, CD14, CD19, CD56 and CD11c), APC-Vio770 HLA-DR, PerCPVio770-CD123 (IL-3R) and PE-CD304 (BDCA4) Abs. The purity of pDC was >98%.

### ELISA assays

PBMC (1-2*10^6 cells) and pDC (1-4*10^4) were maintained for 16 h in RPMI1640+10% FBS+ 1% (PS) (Gibco Laboratories) in 96 well round bottom plates. Cells were cultured with or without 4 *μ*M of TLR7 agonist (Imiquimod) (Sigma-Aldrich), 1 *μ*M of TLR8 or 9 (ORN and ODN2216) (Milteny Biotec) in presence or absence of increasing concentration of CBS004 or control human IgG1 (Crown Biosciences) [0.0005-10 *μ*g/ml]. Cell-free supernatant was harvested after 16 h by centrifugation at 300g for 10 minutes and IFN alpha levels were evaluated using commercially available PBL-Elisa kit (PBL Assay Science), according to manufacturer recommendations. This kit detects 14 out of 15 identified human IFN- α subtypes. They are: IFN- αA, IFN- α2, IFN- αD, IFN- αB2, IFN- αC, IFN- αG, IFN- αH, IFN- αI, IFN- αJ1, IFN- αK, IFN- α1, IFN- α4a, IFN- α4b, and IFN- αWA.

### pDC sub-typing

PBMC were counted and plated in 96 well plate and stimulated for 18 hours using 1 *μ*M ODN 2216. FACS staining was performed as detailed above, using mouse anti-human antibodies directed against lineage markers (VioBlue-CD3, CD14, CD19, CD56 and CD11c), APC-Vio770 HLA-DR, PerCPVio770-CD123 (IL-3R), PE-CD304 (BDCA4), BV650-PD-L1(CD274) and APC-R-700 CD80. Cells then resuspended and analysed using FACS analysis for gating on pDC using Lineage-DR+ CD123+CD304+ and for sub-typing of the three activated pDC populations based on PD-L1 and CD80 expression: P1, PD-L1^+^CD80^−^; P2, PD-L1^+^CD80^+^; P3, PD-L1^−^CD80^+^ (Fig. S1).

### RNA sequencing of healthy pDC

Detailed description of RNA sequencing methods are given in Supplementary Methods file

### Organotypic 3D skin cultures

Primary normal human dermal fibroblasts and keratinocytes (from caucasian female breast tissue) (Promocell) were used to generate a skin-like 3D culture. Firstly, fibroblast-collagen cultures were prepared in Falcon cell culture inserts and placed into Falcon 6 Well Deep Well TC-Treated Polystyrene Plates (BD Biosciences) containing 2×10^5^ fibroblasts in FBS. Once set, 2×10^6^ keratinocytes were seeded on top.

Cultures were placed into Air-Liquid Interphase (ALI) containing Complete KGM Lonza media without BPE supplement, with the addition of 50 **μ**g/ml of ascorbic acid, 1 mg/ml BSA, 10 **μ**g/ml Transferrin, and 1.1 mM of CaCl_2_ (Promocell). Cultures were media changed every 2-3 days and left for 5 days. On day 5, ALI was supplemented with supernatants from pDC treated as above (CTR; no ODN stimulation, ODN; ODN stimulation, ODN+AB; ODN stimulation plus 10 **μ**g/ml CBS004) to produce a final concentration of 6000 pg/ml of IFN in the ODN experiment (determined via ELISA, approx. dilution of supernatants 1:20). Cultures were left for 48 hours. 3 mm punch biopsies were taken and harvested for histology analysis. The remaining culture was collected into 1 ml of TRIzol™ and processed for RNA extraction as described below.

### Xenotransplant mouse models of human pDC activation (XenoSCID)

All mice used were severe combined immunodeficient (CB17/Icr-Prkdcscid/IcrIcoCrl, Charles River) between 4 to 8 weeks of age, housed in accordance with local and Home Office regulations. For the Aldara model, mice were shaved on the back and received topical Aldara application (5% Imiquimod; Meda Health Sales). After 12h, a second application of Aldara and an intraperitoneal (i.p.) injection of CBS004 mAb (5mg/kg) or control human IgG was administrated. 12 h later the mice received an intravenous (i.v.) tail injection of 2.5×10^5^ human pDC. Mice were then euthanized after a further 12h and skin harvested using a punch biopsy and processed for gene expression and FACS analysis. For the bleomycin-induced fibrosis model, 100 **μ**l Bleomycin (Sigma) at 200 **μ**g/ml in PBS was injected subcutaneously into a single location on the shaved back of mice once every other day for 3 weeks. Additionally, some mice received 2.5 ×10^5^ human pDC i.v. on day 0, 7 and 14 following first BLM injection. CBS004 or human IgG (5mg/Kg) were injected i.p. every 5 days starting 24 hours prior to first bleomycin injection.

### ISG response analysis of XenoSCID and organotypic 3D models

RNA was extracted using TRIzol™ Plus RNA Purification Kit (Thermo Fisher Scientific) as per the manufacturer’s instruction. Briefly, skin was homogenized in TRIzol using two 7 mm metal beads and a TissueLyser LT (Qiagen). Homogenates were centrifuged to separate an RNA containing aqueous phase, after which it was further purified by PureLink columns and genomic DNA removed by DNase (Life Technologies, Carlsbad, CA, USA). Eluted RNA was converted to cDNA using RT2 First Strand Kit (Qiagen). Next, the cDNA was mixed with an appropriate RT2 SYBR Green Mastermix (Qiagen). Mouse and Human IFN I RT2 Profiler PCR Arrays (Qiagen) were performed and relative expression determined using the ∆∆CT method and normalized for 5 housekeeping genes according to manufacturer’s guidance.

### Histology

3 mm punch biopsies from mice and patients were formalin-fixed and embedded in paraffin. Sections were cut at 5 **μ**M and subjected to haematoxylin and eosin (H&E) staining. Masson trichrome was used to dye collagen blue and muscle red to identify the extent of fibrosis in the skin samples. For IHC, antigen was retrieved using sodium citrate. Sections were stained with anti-MX1 antibody (Abcam) and pSTAT1 Tyr701 (Cell signaling) followed by ImmPRESS™ (Peroxidase) Polymer Anti-Rabbit IgG Reagent (Vector Laboratories), and visualized with 3, 3-diaminobenzidine (DAB) (Vector Laboratories). Mouse spleen, healthy skin and negative staining were performed for controls. Microscopic analysis was performed using an Olympus BX50 with MicroFire (Optronics) and images captured using Stereo Investigator software at 20X magnification.

### Measurement of collagen in XenoSCID skin samples

Soluble collagen was quantified using the Sircol soluble collagen assay (Biocolor). Punch biopsy skin samples were obtained from XenoSCID and protein extracted and homogenised using M-PER mammalian protein extraction reagent (Thermo Scientific) and two 7mm metal beads. The samples were then further extracted using acetic acid–pepsin solution according to the manufacturer’s protocol.

### FACS analysis of XenoSCID samples

Skin samples from mice were enzymatically digested to release cells using 1 mg/ml collagenase D (Roche), 0.5 mg/ml dispase (Roche) and 0.1 mg/ml DNase-I (Invitrogen, Carlsbad, CA, USA) in Hanks’ balanced salt media (Sigma-Aldrich Corp). For FACS analysis, the released cells were stained with antibodies against human CD45, CD123, CD304 (Miltenyi Biotec). Gating strategy excluded dead cells using Aminoactinomycin D (7-AAD) (BD Biosciences) and sequential gating for human CD45+CD123+ CD304+ as mentioned above.

### Western blotting

Total protein was extracted from skin biopsies in M-PER mammalian protein extraction reagent and resolved by SDS-PAGE (10-15% Tris-Glycine), transferred onto Hybond nitrocellulose membrane (Amersham biosciences) and probed with antibodies specific for MX1. Immunoblots were visualized with species-specific HRP conjugated secondary antibodies (Sigma) and ECL (Thermo/Pierce) on a Biorad ChemiDoc imaging system.

### Statistical analysis

GraphPad Prism 7 software (GraphPad 50 Software) was used for statistical analysis. Pearson’s correlation was used to analyze the association between all studied parameters. One-way analysis of variance combined with Mann-Whitney test or unpaired two tailed t-test were used to evaluate statistically significant differences between groups. Data were expressed as the mean ± standard error (SE). Significance was considered with a P value less than 0.05.

## Acknowledgements

F.D.G., C.S.M., Y.M.E.S., S.H. and R.L.R. designed the study. R.L.R., C.C., G.M., Y.M.E.S. performed the experiments and analysed the data, with additional help from C.W., I.G. and A.C.․ RNAseq analysis was performed by I.C. and A.A.․ J.H.W.D. gave conceptual advice and helped with data interpretation and manuscript draft. R.L.R and F.D.G. wrote the manuscript draft. All authors contributed to draft review.

