## Supplementary material for "Targeting human Plasmacytoid dendritic cells through BDCA2 prevents inflammation and fibrosis in xenotransplant mouse model of Scleroderma": Sup. Materials

### Supplementary Figures

Fig. S1. Gating strategy of pDC within PBMC.

Fig. S2. Production of FAB fragment of CBS004 and dose-response inhibition of IFN alpha secretion.

Fig. S3. Systematic diagram and timeline of *in vivo* experiment of XenoSCID treated with Aldara.

Fig. S4. *In vitro* analysis of IFN secretion by ODN activated purified pDC used in *in vivo* experiments.

Fig. S5. Systematic diagram and timeline of *in vivo* experiment of XenoSCID treated with bleomycin.

### Supplementary Methods

FAB production of CBS004
RNA sequencing and analysis

### Supplementary Tables ( Separate excel files)

STab.1. RNA-seq analysis of Differentially Expressed transcripts ≥ or ≤2-fold change (FDR ≤ 0.05) between CTR and ODN-stimulated pDC.

STab.2. RNA-seq analysis of 87 genes that were increased ≥2-fold (FDR <0.05) between CTR and ODN that are dependent on BDCA2 treatment (reduced ≥1.5 fold by CBS004).

### Supplementary Figures

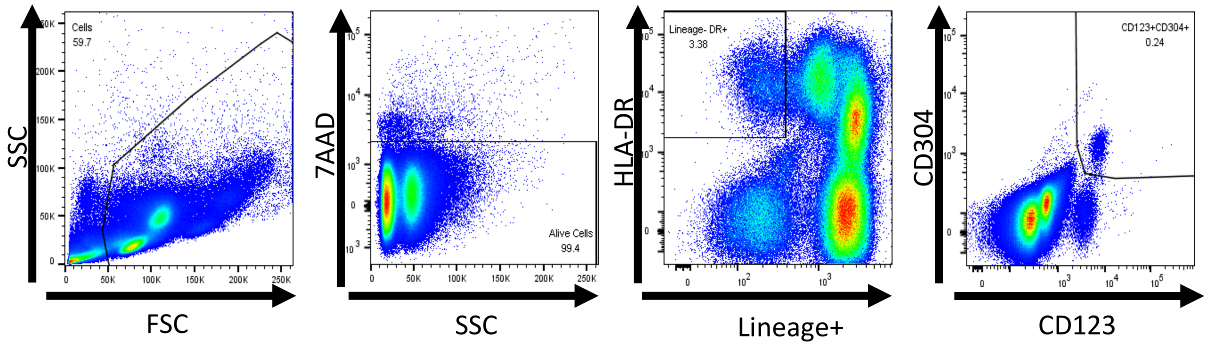


**Fig. S1. Gating strategy of pDC within PBMC.**

PBMC were stained with 7AAD, HLA-DR, Lineage markers (Vioblue-CD3, CD14, CD19, CD56 and CD11c), CD123, CD304, CBS004 and AC144. pDC gated as CD304+ and CD123+ on cells gated as being 7AAD+, HLA-DR+, Lineage- and analysed by flow cytometry. Data from 1 donor shown.

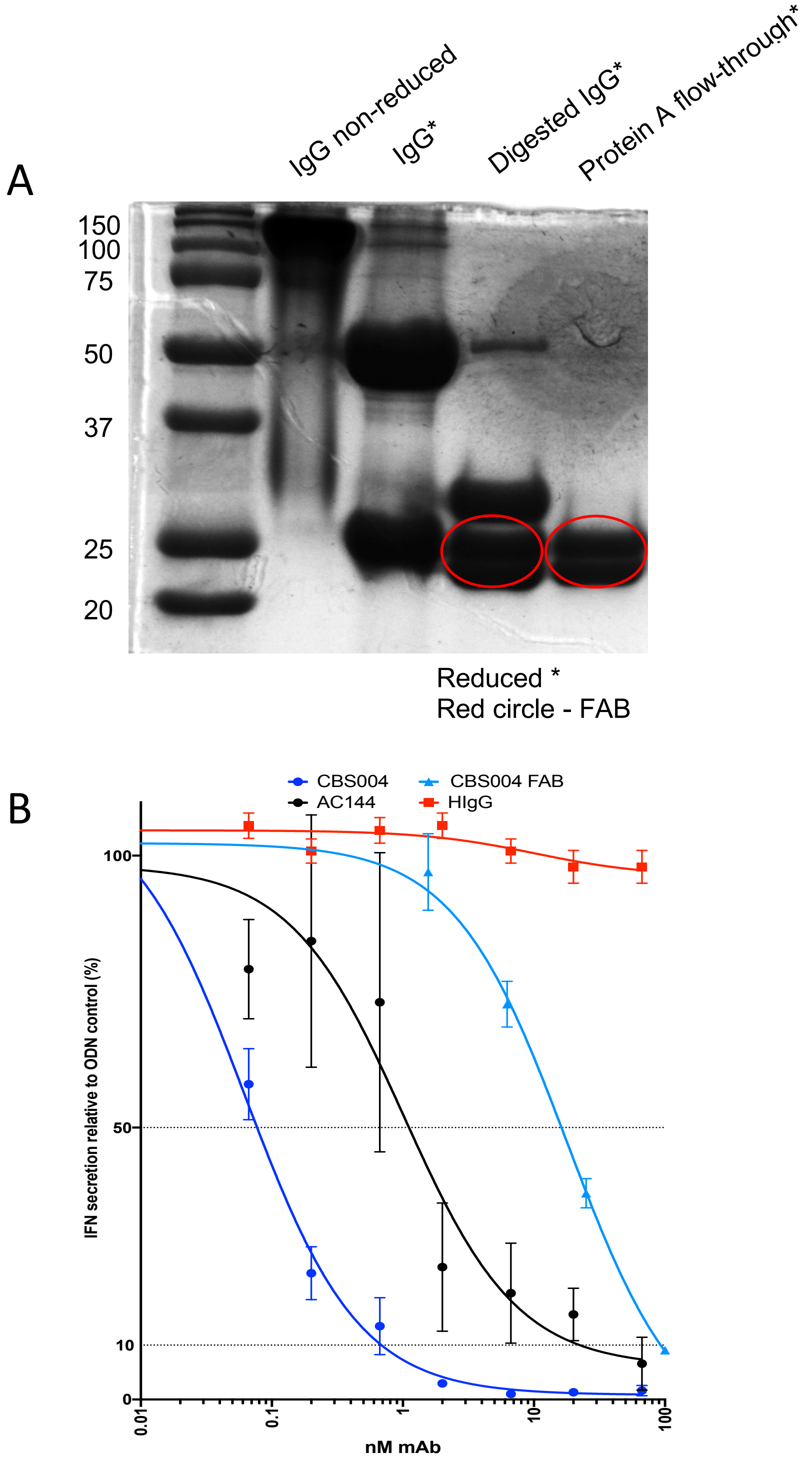


**Fig. S2. Production of FAB fragment of CBS004 and dose-response inhibition of IFN alpha secretion.** (A) SDS-PAGE analysis of undigested IgG, IgG digestion, and elute from Protein A purification column. Non-reduced and reduced conditions were achieved using loading buffers native sample buffer and Laemmli buffer, respectively, and visualized using Coomassie blue. Red circle highlights FAB fragments. (B) Percentage of IFN$\alpha$ secretion, measured by ELISA, from PBMC from 4 donors stimulated with ODN [1 $\mu$M] in the presence of CBS004 FAB, CBS004, AC144 and HIgG [0-100 nM] relative to ODN-stimulated pDC with no antibody. Dotted lines highlight IC50s and IC90s.

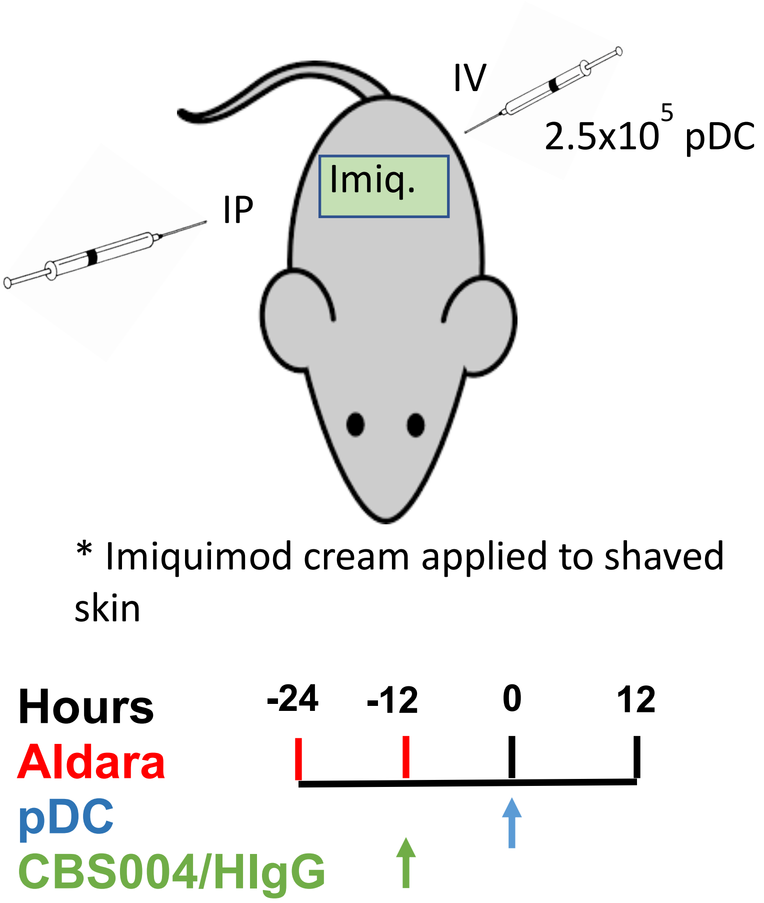


**Fig. S3. Systematic diagram and timeline of *in vivo* experiment of XenoSCID treated with Aldara.** Timeline of *in vivo* experiment, outlining the application of Aldara cream, intravenous (i.v.) tail injection of $2.5x10^5$ human purified pDC and intraperitoneal (i.p.) injection of CBS004 mAb (5 mg/kg) or control human IgG to NODScid mice. Four different treatment conditions consisting of Aldara, Aldara+pDC, Aldara+pDC+CBS004 and Aldara+pDC+HIgG, each in duplicates. Treated skin was harvested using a 3 mm punch biopsy.

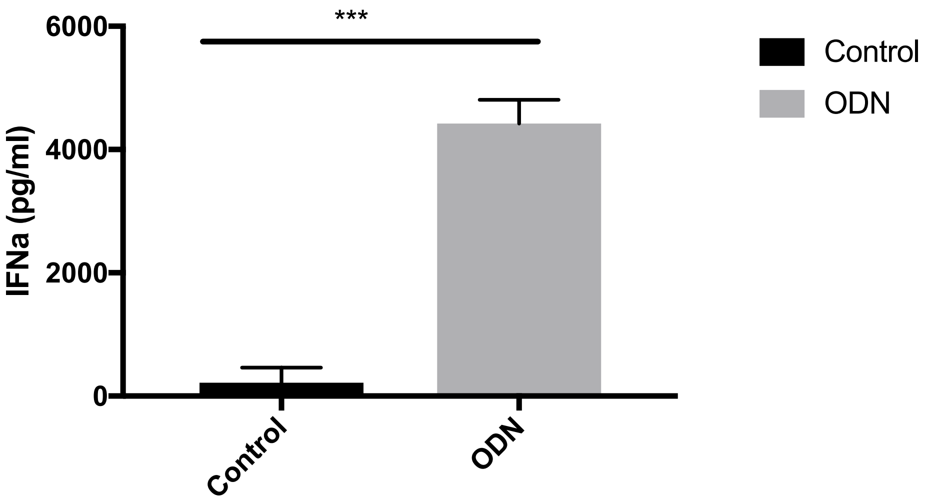


**Fig. S4. *In vitro* analysis of IFN secretion by ODN activated purified pDC used in *in vivo* experiments.** IFNα secretion from purified healthy pDC (n=4) cultured in RPMI alone (CTR), or with ODN (1 μM), after 16 h measured by ELISA. All resulted are represented as means ± SEM. ****P* <0.001 (unpaired two tailed t test).

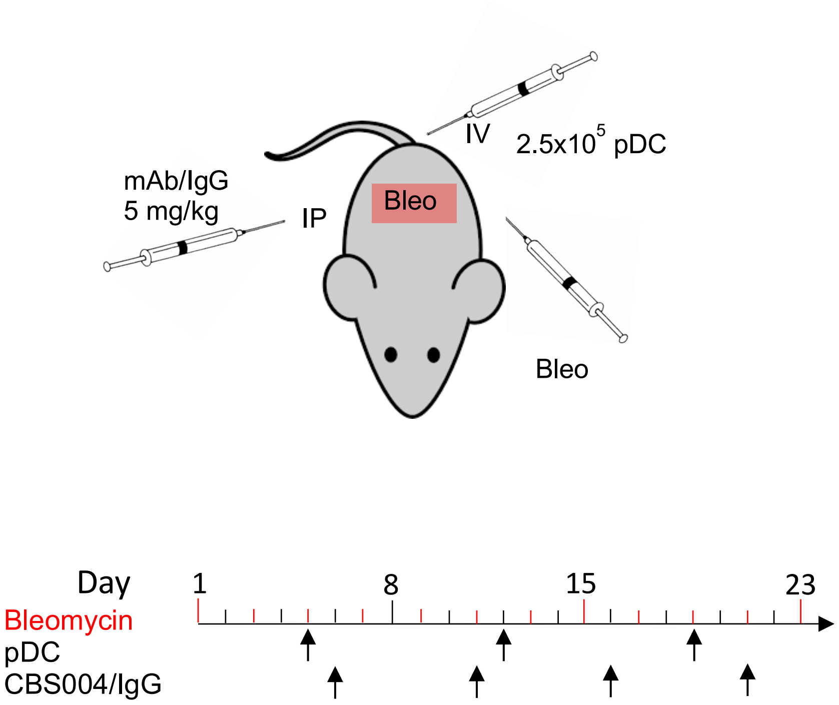


**Fig. S5. Systematic diagram and timeline of *in vivo* experiment of XenoSCID treated with bleomycin.** Diagrams outlining the bleomycin (Bleo) injections, intravenous (i.v.) tail injection of $2.5 \times 10^5$ human purified pDC and intraperitoneal (i.p.) injection of CBS004 mAb (5mg/kg) or control human IgG to NODScid mice. Five different treatment conditions consisting of PBS/Control, Bleo, Bleo+pDC, Bleo+pDC+CBS004 and Bleo+pDC+HIgG, each in triplicates. Treated skin was harvested using a 3 mm punch biopsy.

**Supplementary Methods**

### FAB production of CBS004

Pierce™ Fab Preparation Kit (ThermoFisher Scientific) was used according to manufacturer instructions to generate purified FAB-CBS004. This kit uses papain, a non-specific thiol-endopeptidase, immobilized on agarose resin, to digest IgG to 50kDa Fab and Fc fragments. Digestion was confirmed by running the digest and wash fractions on SDS-PAGE gels using both non- and reducing loading buffers (Native Sample Buffer and Laemmli Buffer (Bio-Rad), respectively) and visualized using Coomassie blue. Digested antibodies were then passed through a NAb Protein A Plus Spin Column to bind Fc fragments and undigested IgG, and purify FAB. Protein concentration of FAB (protein A flow-through and wash) was measured by absorbance at 280 nm and calculated using estimated extinction coefficient of 1.4.

### RNA sequencing and analysis

Total RNA was harvested from commercial healthy human pDC. Firstly, cells were thawed from 4 donors according to manufacturer protocol and cultured as above in RPMI1640+10% FBS+1% PS (unstimulated), 1 $\mu$M ODN2216 with and without 10ug/ml CBS004. RNA was extracted from cells using RNeasy minikit (Qiagen) according to manufacturer's protocol. Ovation® RNA-Seq System V2 (NuGEN) was used to amplify total RNA from all samples. Briefly, first-strand cDNA was made and used to generate double-stranded cDNA followed by a SPIA® amplification. cDNA were quantified by using Qubit dsDNA BR Assay kit (Thermo Fisher Scientific) and the quality was checked by using D1000 screen tape on a Tapestation (Agilent). Covaris S2 sonicator (Woburn) was used to fragment all the cDNA at a size of 200bp. 50 ng cDNA was used to make libraries by using NEBNext® Ultra™ DNA Library Prep Kit for Illumina (Ipswich) without any size selection. The size distribution of the final libraries were checked using the Tapestation and quantified using Quant-iT™ PicoGreen™ dsDNA Assay Kit (Thermo Fisher Scientific). All the libraries were pooled at a concentration of 10 ng and was sequenced on a HiSeq 3000 instrument (Illumina). Pooled sequence data were demultiplexed using Illumina bcl2fastq software, allowing no mismatches in the read index sequences. Raw paired-end sequence data in Fastq format were quality-checked using FastQC software [39]. Cutadapt software [40] was used to trim poor quality bases (Phred quality score < 20) and contaminating adapter sequences from raw reads. Reads trimmed to fewer than 30 nucleotides and orphaned mate-pair reads were discarded. Reads were aligned to human hg38 analysis set reference sequences, obtained from UCSC database [41] using the splicing-aware STAR aligner [42]. STAR aligner was run in 2-pass mode, with known splice junctions supplied in GTF file format, obtained from hg38 RefSeq gene annotation table from UCSC database using Table Browser tool [43]. The resulting alignments in BAM file format were checked for quality using QualiMap software [44] and Picard tools [45], with the latter being used to also mark PCR/Optical duplicate alignments. BAM files were sorted and indexed using Samtools software [46] and visualised using IGV browser [47]. Bioconductor R package RSubread [48] was used to extract raw sequenced fragment counts per transcript using RefSeq hg38 transcript annotation set. Paired-end reads were counted as a single fragment and multi-mapping read pairs were counted as a fraction of all equivalent alignments. Raw count data were normalised for library size differences using median ratio method [49], as implemented in DESeq2 R Bioconductor package [50]. DESeq2 was also used to perform additional data QC steps and differential expression analyses. False Discovery Rate (FDR) was calculated using Benjamini-Hochberg multiple testing correction. Genes below 5% FDR threshold were considered differentially expressed. Differentially expressed gene expression was visualised as clustered heatmaps using the Pheatmap R package [51], using log-transformed normalised gene expression values as input. Principal Component Analysis (PCA) was carried out using the 'prcomp' R function, using the expression of 1000 most variable genes as input. Gene enrichment analyses and annotation were performed using R Bioconductor packages clusterProfiler [52] and ReactomePA [53]. Additionally, KEGG [54] pathways were visualised using Pathview package [55].
